# Genomic Characterisation of Multidrug-Resistant Pathogenic Enteric Bacteria from healthy children in Osun State, Nigeria

**DOI:** 10.1101/2023.07.19.549742

**Authors:** Jessica N. Uwanibe, Idowu B. Olawoye, Christian T. Happi, Onikepe A. Folarin

## Abstract

Antimicrobial resistance (AMR) has been established to be a significant driver for the persistence and spread of bacterial infections. It is, therefore, essential to conduct epidemiological surveillance of AMR in healthy individuals to understand the actual dynamics of AMR in Nigeria. Multi-drug resistant *Klebsiella quasivariicola* (n=1)*, Enterobacter hormaechei* (n=1), and *Escherichia coli* (n=3) from stool samples of healthy children were subjected to whole genome sequencing using Illumina Nextseq1000/2000 and Oxford nanopore. Bioinformatics analysis reveals antimicrobial resistance, virulence genes, and plasmids. This pathogenic enteric bacteria harbored more than three plasmid replicons of either Col and/or Inc type associated with outbreaks and AMR resistant gene *pmrB* responsible for colistin resistance. Plasmid reconstruction revealed an integrated *tetA* gene responsible for tetracycline resistance, and *caa* gene responsible for toxin production in two of the *E.coli* isolates, and a *cusC* gene known to induce neonatal meningitis in the *K. quasivariicola* ST3879. The global spread of MDR pathogenic enteric bacteria is a worrying phenomenon, and close surveillance of healthy individuals, especially children, is strongly recommended to prevent the continuous spread and achieve the elimination and eradication of these infections. Molecular epidemiological surveillance using whole genome sequencing (WGS) will improve the detection of MDR pathogens in Nigeria.

## INTRODUCTION

Enterobacteriaceae are normal inhabitants of the small and large gastrointestinal tracts and are sometimes called enterics. Although most are commensals, some are said to be pathogenic, mainly when found in other areas of the human body (1). Enteric pathogens are a significant cause of morbidity and mortality among young children, especially in low and middle-income countries (LMICs). However, many enteric bacterial infections are either asymptomatic or result in only mild-to-moderate disease (2). Some common enteric pathogens which cause infection in children are *Citrobacter freundii*, *Klebsiella spp*., *Escherichia coli*, *Enterobacter spp, Salmonella typhi, and Shigella spp* (3, 4).

Although the exact burden of infectious diseases caused by enteric bacteria in Sub-Saharan Africa is vague, recent reports have shown an increase in the prevalence of diseases caused by enteric bacteria, such as sepsis and typhoid (5), cholera (6), shigellosis (7), and diarrhoeal infection (8). Antimicrobial resistance (AMR) has been established to be a significant driver for the persistence and spread of these diseases, particularly in cases where the causative agents are multidrug-resistant (MDR) or extensively drug-resistant (XDR) (9–11). The overuse and misuse of antibiotics has been identified as a leading cause of AMR in enteric bacteria, as it creates selective pressure and promotes the emergence of resistance, leading to an alarming increase in the prevalence of MDR enteric bacteria among healthy individuals (12). However, the ease of transfer of plasmids and transposons harboring genes associated with AMR, amongst enterics, through horizontal gene transfer ensures the persistence and spread of AMR (13–15).

In Nigeria, AMR is still a significant public health concern. The prevalence has been reported to range from 31.14% to 90.5% in North and Southern Nigeria (16, 17). Despite government interventions and policies, there needs to be more effective epidemiological surveillance and database for AMR in enteric bacteria of public health importance in the country. Furthermore, there are few in-depth genomic analyses of enteric bacteria of public health interest. (18, 19).

Genomic surveillance of enteric bacteria species of public health importance, such as Escherichia coli, *Klebsiella pneumoniae*, *Salmonella typhi*, *Vibrio cholera*, and *Shigella spp* not only informs treatment guidelines for diseases caused by these human pathogens but is necessary for the design and implementation of AMR interventions and control measures (20). Whole genome sequencing (WGS) provides a high-resolution method capable of providing detailed characterization and epidemiological information on pathogen genomic diversity, transmission, and evolution, as well as in-depth genomic data on AMR dynamics and spread, aiding in developing better control measures (21).

A healthy human microbiome is one of the major reservoirs of antibiotic-resistant genes (ARGs), which are transmissible to pathogenic bacteria (22). Recent research reveals that antibiotic-resistant genes (ARGs) contribute to the prevalence of multidrug-resistant (MDR) enteric bacteria in healthy humans, ensuring the circulation of ARGs in the population(16). Children are the major population bearing this burden, with cases of child mortality due to diarrheal disease caused by resistant pathogenic enterics on the rise (23, 24). Therefore, it is essential to conduct epidemiological surveillance of AMR in healthy individuals to understand the actual dynamics of AMR in Nigeria fully. This will inform policy that will drive the achievement of elimination and eradication of diseases.

This study aimed to characterize AMR genes and mobile genetic elements harbored in the multidrug-resistant Enterobacteriaceae isolates from healthy children.

## METHODS

### Ethical approval

This study was conducted according to the guidelines of the Declaration of Helsinki and was approved by the Ethics Committee from the Research Ethics Committee of the Ladoke Akintola University of Technology Teaching Hospital (LAUTECH) (LTH/EC/2019/09/431) and the Ministry of Health (OSHREC/PRS/569T/164) Oshogbo, Osun State, Nigeria. Informed consent was obtained from all subjects involved in this study.

### Bacteria Isolation, Identification and Selection

A total of 5 enteric isolates (*E. coli* [*n* = 3]; *Enterobacter spp* [*n* = 1]; *Klebsiella pneumonia* [*n* = 1]) collected between 2019 to 2020 were recovered from a previous study carried out in Osun state for which a serological profile has been published (25). These multidrug-resistant isolates were obtained from stool samples from presumptively healthy children under the age of 15 years. Samples were cultured on Salmonella/Shigella and MacConkey agar and Initial identification was performed with API 20E test kit (BioMeriuxe, Marcy-l’Étoile, France).

### Antimicrobial Susceptibility Test

Antimicrobial susceptibility testing was performed using the disk diffusion technique. The antibiotics disc used were; Gentamicin (30 µg), Tetracycline (30 µg), Ciprofloxacin (5 µg), Chloramphenicol (30 µg), Trimethoprim/sulfamethoxazole (25 µg), Ceftazidime (30 µg), Ceftriaxone (30 µg), Cefotaxime (30µg) and Aztreonam (30 µg) as described previously by Kariuki and others (26). Results were interpreted according to the 2017 guidelines provided by the Clinical and Laboratory Standards Institute (CLSI). Results were analyzed and interpreted using the ABIS online software (http://www.tgw1916.net/bacteria_logare_desktop.html) (17).

### Whole Genome Sequencing

All isolates were subcultured before DNA extraction. DNA was extracted using the Qiagen DNeasy Blood and Tissue kit (Qiagen, USA). Extracted samples were quantified using a Qubit fluorometer (ThermoFisher Scientific) using a dsDNA high-sensitivity assay. Sequencing libraries were prepared using the Nextera DNA flex preparation kit (Illumina, USA). Library preparation protocol was adopted from the CDC PulseNet Nextera DNA Flex Standard operating protocol and sequenced using the Illumina Miseq platform and Nextseq 1000/2000 at the African Center of Excellence for Genomics of Infectious Diseases (ACEGID), Redeemer’s University, Nigeria.

To improve the plasmid assembly, we performed a single run on a GridION x5 to generate long reads. Library preparation and sequencing were done using the Rapid PCR Barcoding kit (SQK-RPB004) (Oxford Nanopore Technologies, Oxford, UK), following the manufacturer’s recommendations. We used a GridION MK1 sequencer, FLO-MIN106D R9 flow cell, and MinKNOW software v22.10.7 for sequencing.

### Genomic data analysis

Raw FASTQ files were processed with the Connecticut Public Health Laboratory (CT-PHL) pipeline, also known as C-BIRD v0.9 (https://github.com/Kincekara/C-BIRD). The assembled contigs were further checked for contamination using CheckM(27). Isolates with a bracken taxon ratio < 0.7, genome estimated ratio > 1.1, estimated sequencing depth < 40x, and genome completeness < 90% were excluded from subsequent analyses. Further analyses, including genome annotation, plasmid detection, antimicrobial resistance, and virulence prediction, and MLST typing, were performed with the Public Health Bacterial Genomics (PHBG) v1.3.0 (https://github.com/theiagen/public_health_bacterial_genomics). Genome assemblies of isolates were refined with the unicycler (28) hybrid assembler using Illumina and Oxford nanopore sequence reads. The plasmids were reconstructed and typed with MOB-suite (29) and annotated with pLannotate (30) and Prokka (31).

### Data availability

Sequence data of isolates are deposited in NCBI Sequence Read Archive (SRA) under BioProject accession number PRJNA838568

## RESULT

### Antibiotic susceptibility test and AMR prediction

All isolates subjected to antibiotic susceptibility tests were multidrug-resistant. All the isolates were resistant to only ciprofloxacin and either/or cefotaxime, ceftriaxone, ceftazidime, tetracycline, chloramphenicol, gentamicin and sulfamethoxazole-trimethoprim (Table 1).

**Table 1.** Antimicrobial resistance profile of the five (5) enteric isolates from healthy children.

| Drug Class | Antibiotics | Susceptible<br>N (%) | Intermediate<br>N (%) | Resistance<br>N (%) |
| --- | --- | --- | --- | --- |
| Beta-lactam<br>(Cephalosporines) | Cefotaxime | 1/(20%) | 0 | 4(80%) |
|  | Ceftriaxone | 0 | 3(60%) | 2(40%) |
|  | Ceftazidime | 2(40%) | 0 | 3(60%) |
| Tetracycline | Tetracycline | 1(20%) | 1(20%) | 3(60%) |
| Quinolones | Ciprofloxacin | 0 | 0 | 5(100%) |
| Phenolics | Chloramphenicol | 1(20%) | 1(20%) | 3(60%) |
| Aminoglycosides | Gentamicin | 0 | 1(20%) | 4(80%) |
| Folate pathway | Trimethoprim/Sulfamethoxazole | 0 | 1(20%) | 4(80%) |

N= Number of isolates.

### Genome sequence analysis of the isolates

Genome assembly analysis shows a genome length of 4.8 Mbp to 6 Mbp, 5.6 Mbp, and 5.8 Mbp for *E. coli*, *K. quasiveriicola,* and *Enterobacter hormaechei,* respectively. Through probing, the assembled contigs of the bacteria genomes with antimicrobial resistance (AMR) detection software, such as AMRFinder+ and Kleborate, showed the presence of beta-lactams resistance genes such as *bla*_TEM-1_, *bla*_EC_, *bla*_ACT_, and *bla*_OKP-D_, (Table 1). In addition to this, all five isolates also contained other AMR genes associated with either fluoroquinolone (*gyrAS83L, qnrS1*), aminoglycoside (*aph*(*6*)*-Id, aph(3’’)-Ib*), tetracycline (*tetA*), and sulfonamide (*sul1, sul2*) resistance (Table 2). MLST typing predicted ST type for three isolates, *E.coli* ST 219, ST450, and *K. quasiveriicola* ST 3897, while the remaining two isolates, *Enterobacter hormaechei,* and *E. coli,* had no ST prediction.

**Table 2.** AMR genes detected from Sequenced MDR Enteric Bacteria (*n* = 5).

| Drug class | AMR genes (number of isolates) |
| --- | --- |
| Beta-lactam (Cephalosporines) | *BlaTEM-1* (3)*, *BlaOXA-1* (1)*, *BlaEC* (3)*, *BlaOKP-D* (1)*, *BlaACT-16* (1)*. |
| Efflux pumps | *pmrB_Y358N* (1)*, *acrF* (3)*, *oqxB* (2)*, *oqxA* (2)*, *qepA4* (1)*, *mdtM* (3)*, *emrD* (2)*. |
| Tetracycline | *tetA* (3)*. |
| Quinolones | *gyrA_S83L* (2)*, *qnrS1* (1)*, *gyrA_D87N* (1)*, *parC_S80I* (1)*, *parE_S458A* (1)*. |
| Phenolics | *catA1* (1)*. |
| Aminoglycosides | *aph(6)-Id* (2)*, *aph(3'')-Ib* (2)*, *aadA1* (1)*, |
| Fosfomycin | *fosA* (2)*, *glpT_E448K* (1)*. |
| Folate pathway | *Sul1* (1)*, *Sul2* (2)*, *dfrA7* (1)*, *dfrA17* (1)*. |

\*indicate the number of isolates

### Analysis for Virulence genes and Mobile genetic elements in the isolates

Virulence genes were present in all the 3 *E.coli* isolates. At the same time, none was detected in *K. quasivariicola* and *Enterobacter hormaechei.* Virulence genes responsible for bacteria iron uptake were detected in all the *E.coli* isolates. Aerobactin virulence genes, *iucABCD,* and *iutA* were also present in all *E.coli* isolates, but only two *E.coli* isolates harbored the Yersiniabactin virulence genes *ybtP* and *ybtQ*. Other virulence genes detected include gene encoding enterotoxin *senB,* increased serum survival *iss* gene, *sigA,* secreted autotransporter toxin -*sat,* east-1 heat-stable toxin -*astA,* long polar fimbriae -*ipfA, LEE* encoded type III secretion system effector-*espX1,* for adherence-*fdeC,* P-fimbria operon-*papHCFGII,* Salmonella HilA homolog-*eilA,* and iron-regulated outer membrane virulence protein - *ireA* gene.

A total of 28 plasmids was detected in these isolates, harboring between 3 and 9 plasmid replicons associated with antibiotics resistance genes (ARGs). The Inc plasmid type is the most occurring plasmid across all isolates. IncF and Col plasmids were detected in 90% (4/5) of the isolates. IncF replicons detected include IncFII(pBK30683) (1/5); InFIB(K) (2/5); IncFII(K) (1/5); IncFII(pECLA) (1/5); IncFIA(HI1) (1/5); IncFIA (1/5); IncFIB(AP001918) (1/5); and IncFII(pRSB107) (1/5) while Col plasmid had replicons which included Col440I (2/5); Col440II (1/5); ColpHAD28 (3/5); ColMG828 (1/5) and Col156 (1/5). Other plasmid replicons detected were, IncR (1/5), IncI2 (1/5), IncQ1(1/5), and IncB/O/K/Z (2/5).

#### PLASMID RECONSTRUCTION USING OXFORD NANOPORE SEQUENCING

We reconstructed 4 plasmids ranging from 3,536 to 163,036 bp detected in the *K. quasivariicola* isolate, and were either conjugative, non-mobilizable, or mobilizable plasmids. The largest plasmid contained 11 contigs with IncFIB, IncFII, rep_cluster_2183, rep_cluster_2327, and rep_cluster_2358 replicon types. None of the plasmids harbored any antibiotic-resistant gene however, several metal-resistant genes and efflux proteins such as *copA*, *copB*, *silP*, *cusA*, *cusB*, and *cusC* were integrated into the plasmid (Fig. 1). We also detected the tetracycline resistance gene *tetA* and another *caa* gene integrated into the plasmid of two *E. coli* isolates that codes for colicin polypeptide toxins. (Table S1).

**Figure 1.**
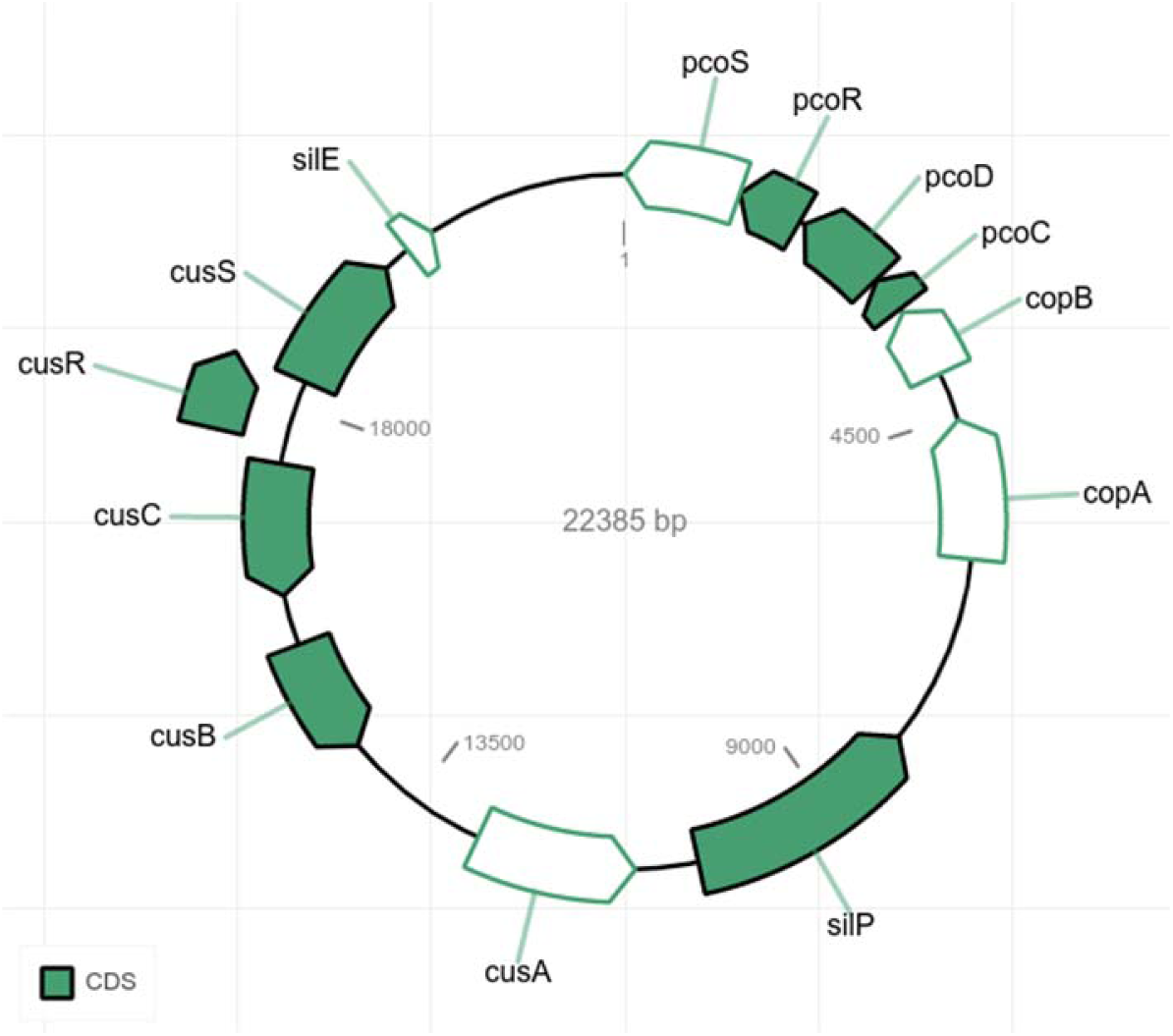
Coding sequences of one of the contigs of the pKPC-0cc9 plasmid detected in the *K. pneumoniae* isolate. Solid green-colored CDS represents > 90% match length, while white-colored CDS represents < 90% match length of the query.

## DISCUSSION

Antimicrobial resistance, especially in enteric bacteria of public health interest, continues to be of global concern, especially in low and middle-income countries (17). In Nigeria, only a handful of studies provide a genomic characterization of circulating multidrug-resistant enteric bacteria, especially in healthy individuals that serve as reservoirs. Therefore, our study aimed to use genomic tools to characterize multidrug-resistance enteric bacteria isolated from healthy children in a semi-urban area of Osun State, Nigeria. In this study, we recovered enteric pathogens of public health interest, such as *Klebsiella quasivariicola*, *Enterobacter hormaechei,* and *Escherichia coli*. The occurrence of these pathogenic enteric bacteria in febrile individuals is well-documented across the country(17, 32, 33); however, little or no information on healthy individuals.

This study revealed the presence of beta-lactam-resistant genes in all five isolates, fluoroquinolones resistant genes with more than triple mutation (*gyrA_S83L, gyrA_D87N, parC_S80I,* and *parE_S458A*), the Colistin-resistant gene (*pmrB_Y358N*) found in XDR E.coli (34, 35) and other classes of antibiotics. The presence of these enteric pathogens isolated from presumptive healthy children is of great concern as these children serve as potential reservoirs for transmission of these pathogens to other susceptible individuals. In addition to transmission, the presence of the resistance genes indicates the possibility of treatment failure, thus promoting longer existence in the population. It is possible that these pathogens developed resistance due to consistent drug pressure, as most drugs are readily available to the citizens in the country.

Virulence genes and plasmid replicons associated with XDR carriage were also identified(35). This provides insight into the genetic variation of pathogenic enterics in healthy children in the southwestern part of Nigeria. Virulence genes identified amongst the *E. coli* isolates are known to play a significant role in influencing the degree of pathogenicity of bacteria infection in confirmed disease patients in some reported cases (36, 37). Although the *K. quasivariicola* and *Enterobacter hormaechei* harbored no virulence gene, the presence of more than three plasmid replicons associated with ARGs is a significant concern as reports of colonizing enteric bacteria resulting in infections are facilitated by MDR and plasmid carriage (38).

The pathogenic enteric bacteria in this study are known as commensals of the gut microbiota and have been reported in other studies (39–41). Yet, the presence of MDR genes and plasmids harbored by these bacteria in healthy children is of concern as children in this community will enable the persistent transmission of diseases caused by these enteric bacteria leading to an increase in community-acquired infections(42). This could increase morbidity, mortality, and healthcare cost, especially in immunocompromised individuals.

The detection of AMR genes in this population of healthy children, although not surprising, but alarming, as antibiotics abuse has been established in this part of the world (18, 43). This is seen in the AST result and genomic data, as most isolates were resistant to folate-pathway drugs, quinolones, and beta-lactam drugs. The presence of efflux pumps responsible for MDR resistance increases the expression of resistance in these isolates. Resistance to these drugs, some of the most common antibiotics used in treating diarrhea diseases in children, could pose a challenge in future infections with these resistant strains. Also, reports have shown that most cases of infection with MDR enterics are not just from a person-to-person transmission or from contaminated water/food but could also be from the individual colonized by these MDR bacteria (38).

A significant driver of the spread of AMR genes in bacteria is the presence of mobile genetic elements (44). Plasmids spread AMR genes to other bacteria through horizontal gene transfer, thereby increasing the prevalence of antibiotic resistance. Plasmids are known to contain genes responsible for antibiotics resistance, colonization, and virulence, which provides an advantage for bacteria survival(44); as seen in this study, the plasmid recovered from one of the *E. coli* strains harbored a *tetA* gene responsible for tetracycline resistance and *caa* gene that codes for colicin polypeptide toxins known to destroy cells of other organisms by depolarizing the cytoplasmic membrane of the cells (45, 46). Also, the plasmid in the *K. quasivariicola* harbored the *cusC* gene associated with copper and silver resistance, which, when found in pathogenic *K. pneumoniae,* facilitates the invasion of the brain microvascular endothelial cells, thereby causing neonatal meningitis (47, 48).

Apart from the presence of virulence genes and mobile genetic elements as a driver of MDR, the indiscriminate use of antibiotics in this part of the world is also a significant contributing factor to the high prevalence of MDR bacteria (19, 49). Although the healthy children in this study were not on any antibiotics drugs at the time of sample collection, it has been reported that children are exposed to antibiotics early on, mainly without a physician’s prescription (50–52). To mitigate the spread of MDR enteric bacteria and the possible emergence of XDR enteric bacteria, there is a need for proper surveillance in healthy individuals to achieve proper monitoring and control of AMR in the country. Although some control measures have been implemented to monitor AMRs in the population, the neglect in surveying healthy individuals will truncate all efforts. Another key factor in proper surveillance is the use of sensitive techniques. Whole genome sequencing employed in this study revealed genetic elements driving pathogenicity and the spread of MDR enteric bacteria from healthy children. This might not have been possible by culture or PCR alone. This proves the importance of the WGS technique in pathogen enteric bacteria surveillance and its implantation also in healthy individuals to be able to achieve the one health approach in combating disease eradication.

## CONCLUSION

Our findings showed that whole genome sequencing and its in-depth analysis could accurately reconstruct the molecular characterization of isolates and advance the interpretation of bacteria. The global spread of MDR enteric bacteria in healthy children is a worrying phenomenon, and close inspection to avoid their spread is strongly needed. Applying WGS in surveillance will improve the detection of MDR pathogens by overcoming the limitation of analyzing only a small part of the genome and providing more rapid management, and controlling the emergence of new antibiotic-resistant strains and their evolution. Since the human gut microbiome is a hotspot for antibiotic resistance exchange and evolution, it is recommended to perform active and continuous surveillance of these bacteria and their phenotypes to monitor any emergence of novel resistant patterns and carriage of genes for virulence factors and prevalent clones.

## Acknowledgments

We thank the Ladoke Akintola teaching hospital for their collaborative support in conducting the study. We are also grateful for the facility at the African Center of Excellence for Genomics of Infectious Diseases (ACEGID), Redeemer’s University, Nigeria, that enabled us to carry out this study.

## Contribution to Authorship

Study concept and design: JNU, OAF ; sequencing and bioinformatics: JNU and IBO; analysis and interpretation of data: JNU and IBO authors; funding acquisition: TCH; drafting of the manuscript: JNU and IBO critical revision of the manuscript for important intellectual content: JNU, IBO, TCH and OAF.

## REFERENCE

1. Rock C, Donnenberg MS. 2014. Human Pathogenic EnterobacteriaceaeReference Module in Biomedical Sciences. Elsevier.

2. Lee GO, Eisenberg JNS, Uruchima J, Vasco G, Smith SM, Van Engen A, Victor C, Reynolds E, MacKay R, Jesser KJ, Castro N, Calvopiña M, Konstantinidis KT, Cevallos W, Trueba G, Levy K. 2021. Gut microbiome, enteric infections and child growth across a rural–urban gradient: protocol for the ECoMiD prospective cohort study. BMJ Open 11:e046241.

3. Fletcher SM, Stark D, Ellis J. 2011. Prevalence of gastrointestinal pathogens in Sub-Saharan Africa: systematic review and meta-analysis. J Public Health Africa 2:e30.

4. Oppong TB, Yang H, Amponsem-Boateng C, Kyere EKD, Abdulai T, Duan G, Opolot G. 2020. Enteric pathogens associated with gastroenteritis among children under 5 years in sub-Saharan Africa: a systematic review and meta-analysis. Epidemiol Infect 148:e64.

5. Kim CL, Cruz Espinoza LM, Vannice KS, Tadesse BT, Owusu-Dabo E, Rakotozandrindrainy R, Jani IV, Teferi M, Bassiahi Soura A, Lunguya O, Steele AD, Marks F. 2022. The Burden of Typhoid Fever in Sub-Saharan Africa: A Perspective. Res Rep Trop Med 13:1–9.

6. Elimian KO, Musah A, Mezue S, Oyebanji O, Yennan S, Jinadu A, Williams N, Ogunleye A, Fall IS, Yao M, Eteng W-E, Abok P, Popoola M, Chukwuji M, Omar LH, Ekeng E, Balde T, Mamadu I, Adeyemo A, Namara G, Okudo I, Alemu W, Peter C, Ihekweazu C. 2019. Descriptive epidemiology of cholera outbreak in Nigeria, January-November, 2018: implications for the global roadmap strategy. BMC Public Health 19:1264.

7. Rogawski McQuade ET, Shaheen F, Kabir F, Rizvi A, Platts-Mills JA, Aziz F, Kalam A, Qureshi S, Elwood S, Liu J, Lima AAM, Kang G, Bessong P, Samie A, Haque R, Mduma ER, Kosek MN, Shrestha S, Leite JP, Bodhidatta L, Page N, Kiwelu I, Shakoor S, Turab A, Soofi SB, Ahmed T, Houpt ER, Bhutta Z, Iqbal NT. 2020. Epidemiology of Shigella infections and diarrhea in the first two years of life using culture-independent diagnostics in 8 low-resource settings. PLoS Negl Trop Dis 14:e0008536.

8. Mero S, Timonen S, Lääveri T, Løfberg S, Kirveskari J, Ursing J, Rombo L, Kofoed P-E, Kantele A. 2021. Prevalence of diarrhoeal pathogens among children under five years of age with and without diarrhoea in Guinea-Bissau. PLoS Negl Trop Dis 15:e0009709.

9. Kamel NA, El-Tayeb WN, El-Ansary MR, Mansour MT, Aboshanab KM. 2019. XDR-Klebsiella pneumoniae isolates harboring blaOXA-48: In vitro and in vivo evaluation using a murine thigh-infection model. Exp Biol Med 244:1658–1664.

10. Ramsamy Y, Mlisana KP, Amoako DG, Allam M, Ismail A, Singh R, Abia ALK, Essack SY. 2020. Pathogenomic Analysis of a Novel Extensively Drug-Resistant Citrobacter freundii Isolate Carrying a blaNDM-1 Carbapenemase in South Africa. Pathogens 9.

11. Zakir M, Khan M, Umar MI, Murtaza G, Ashraf M, Shamim S. 2021. Emerging Trends of Multidrug-Resistant (MDR) and Extensively Drug-Resistant (XDR) Salmonella Typhi in a Tertiary Care Hospital of Lahore, Pakistan. Microorganisms 9.

12. van Duin D, Paterson DL. 2020. Multidrug-Resistant Bacteria in the Community: An Update. Infect Dis Clin North Am 34:709–722.

13. Nikaido H. 2009. Multidrug resistance in bacteria. Annu Rev Biochem 78:119–146.

14. Sun D, Jeannot K, Xiao Y, Knapp CW. 2019. Editorial: Horizontal Gene Transfer Mediated Bacterial Antibiotic Resistance. Front Microbiol 10:1933.

15. Wallace MJ, Fishbein SRS, Dantas G. 2020. Antimicrobial resistance in enteric bacteria: current state and next-generation solutions. Gut Microbes 12:1799654.

16. Aworh MK, Kwaga J, Okolocha E, Mba N, Thakur S. 2019. Prevalence and risk factors for multi-drug resistant Escherichia coli among poultry workers in the Federal Capital Territory, Abuja, Nigeria. PLoS One 14:e0225379.

17. Kayode A, Okunrounmu P, Olagbende A, Adedokun O, Hassan A-W, Atilola G. 2020. High prevalence of multiple drug resistant enteric bacteria: Evidence from a teaching hospital in Southwest Nigeria. J Infect Public Health 13:651–656.

18. Achi CR, Ayobami O, Mark G, Egwuenu A, Ogbolu D, Kabir J. 2021. Operationalising One Health in Nigeria: Reflections From a High-Level Expert Panel Discussion Commemorating the 2020 World Antibiotics Awareness Week. Front Public Health 9:673504.

19. Chukwu EE, Oladele DA, Enwuru CA, Gogwan PL, Abuh D, Audu RA, Ogunsola FT. 2021. Antimicrobial resistance awareness and antibiotic prescribing behavior among healthcare workers in Nigeria: a national survey. BMC Infect Dis 21:22.

20. Afolayan AO, Oaikhena AO, Aboderin AO, Olabisi OF, Amupitan AA, Abiri OV, Ogunleye VO, Odih EE, Adeyemo AT, Adeyemo AT, Obadare TO, Abrudan M, Argimón S, David S, Kekre M, Underwood A, Egwuenu A, Ihekweazu C, Aanensen DM, Okeke IN, NIHR Global Health Research Unit on Genomic Surveillance of Antimicrobial Resistance. 2021. Clones and Clusters of Antimicrobial-Resistant Klebsiella From Southwestern Nigeria. Clin Infect Dis 73:S308–S315.

21. Bolourchi N, Giske CG, Nematzadeh S, Mirzaie A, Abhari SS, Solgi H, Badmasti F. 2022. Comparative resistome and virulome analysis of clinical NDM-1–producing carbapenem-resistant Enterobacter cloacae complex. Journal of Global Antimicrobial Resistance 28:254–263.

22. Kumar M, Sarma DK, Shubham S, Kumawat M, Verma V, Nina PB, Jp D, Kumar S, Singh B, Tiwari RR. 2021. Futuristic Non-antibiotic Therapies to Combat Antibiotic Resistance: A Review. Front Microbiol 12:609459.

23. Okeke IN, Aboderin OA, Byarugaba DK, Ojo KK, Opintan JA. 2007. Growing problem of multidrug-resistant enteric pathogens in Africa. Emerg Infect Dis 13:1640–1646.

24. Mokomane M, Kasvosve I, de Melo E, Pernica JM, Goldfarb DM. 2018. The global problem of childhood diarrhoeal diseases: emerging strategies in prevention and management. Ther Adv Infect Dis 5:29–43.

25. Uwanibe JN, Kayode TA, Oluniyi PE, Akano K, Olawoye IB, Ugwu CA, Happi CT, Folarin OA. 2023. The Prevalence of Undiagnosed Salmonella enterica Serovar Typhi in Healthy School-Aged Children in Osun State, Nigeria. Pathogens 12.

26. Kariuki S, Dyson ZA, Mbae C, Ngetich R, Kavai SM, Wairimu C, Anyona S, Gitau N, Onsare RS, Ongandi B, Duchene S, Ali M, Clemens JD, Holt KE, Dougan G. 2021. Multiple introductions of multidrug-resistant typhoid associated with acute infection and asymptomatic carriage, Kenya. Elife 10.

27. Parks DH, Imelfort M, Skennerton CT, Hugenholtz P, Tyson GW. 2015. CheckM: assessing the quality of microbial genomes recovered from isolates, single cells, and metagenomes. Genome Res 25:1043–1055.

28. Wick RR, Judd LM, Gorrie CL, Holt KE. 2017. Unicycler: Resolving bacterial genome assemblies from short and long sequencing reads. PLoS Comput Biol 13:e1005595.

29. Robertson J, Nash JHE. 2018. MOB-suite: software tools for clustering, reconstruction and typing of plasmids from draft assemblies. Microb Genom 4.

30. McGuffie MJ, Barrick JE. 2021. pLannotate: engineered plasmid annotation. Nucleic Acids Res 49:W516–W522.

31. Seemann T. 2014. Prokka: rapid prokaryotic genome annotation. Bioinformatics 30:2068– 2069.

32. Oyejobi GK, Sule WF, Akinde SB, Khan FM, Ogolla F. 2022. Multidrug-resistant enteric bacteria in Nigeria and potential use of bacteriophages as biocontrol. Sci Total Environ 824:153842.

33. Obasi AI, Ugoji EO, Nwachukwu SU. 2019. Incidence and molecular characterization of multidrug resistance in Gram-negative bacteria of clinical importance from pharmaceutical wastewaters in South-western Nigeria. Environmental DNA 1:268–280.

34. Wang M, Wang W, Niu Y, Liu T, Li L, Zhang M, Li Z, Su W, Liu F, Zhang X, Xu H. 2020. A Clinical Extensively-Drug Resistant (XDR) Escherichia coli and Role of Its β-Lactamase Genes. Front Microbiol 1:590357.

35. Al-Mustapha AI, Raufu IA, Ogundijo OA, Odetokun IA, Tiwari A, Brouwer MSM, Adetunji V, Heikinheimo A. 2023. Antibiotic resistance genes, mobile elements, virulence genes, and phages in cultivated ESBL-producing Escherichia coli of poultry origin in Kwara State, North Central Nigeria. Int J Food Microbiol 389:110086.

36. Sora VM, Meroni G, Martino PA, Soggiu A, Bonizzi L, Zecconi A. 2021. Extraintestinal Pathogenic Escherichia coli: Virulence Factors and Antibiotic Resistance. Pathogens 10.

37. Duan Y, Gao H, Zheng L, Liu S, Cao Y, Zhu S, Wu Z, Ren H, Mao D, Luo Y. 2020. Antibiotic Resistance and Virulence of Extraintestinal Pathogenic Escherichia coli (ExPEC) Vary According to Molecular Types. Front Microbiol 11:598305.

38. Campos-Madueno EI, Moradi M, Eddoubaji Y, Shahi F, Moradi S, Bernasconi OJ, Moser AI, Endimiani A. 2023. Intestinal colonization with multidrug-resistant Enterobacterales: screening, epidemiology, clinical impact, and strategies to decolonize carriers. Eur J Clin Microbiol Infect Dis 42:229–254.

39. Houghteling PD, Walker WA. 2015. Why is initial bacterial colonization of the intestine important to infants’ and children’s health? J Pediatr Gastroenterol Nutr 60:294–307.

40. Oriá RB, Murray-Kolb LE, Scharf RJ, Pendergast LL, Lang DR, Kolling GL, Guerrant RL. 2016. Early-life enteric infections: relation between chronic systemic inflammation and poor cognition in children. Nutr Rev 74:374–386.

41. Milani Christian, Duranti Sabrina, Bottacini Francesca, Casey Eoghan, Turroni Francesca, Mahony Jennifer, Belzer Clara, Delgado Palacio Susana, Arboleya Montes Silvia, Mancabelli Leonardo, Lugli Gabriele Andrea, Rodriguez Juan Miguel, Bode Lars, de Vos Willem, Gueimonde Miguel, Margolles Abelardo, van Sinderen Douwe, Ventura Marco. 2017. The First Microbial Colonizers of the Human Gut: Composition, Activities, and Health Implications of the Infant Gut Microbiota. Microbiol Mol Biol Rev 81:10.1128/mmbr.00036–17.

42. van Duin D, Paterson DL. 2016. Multidrug-Resistant Bacteria in the Community: Trends and Lessons Learned. Infect Dis Clin North Am 30:377–390.

43. Chukwu EE, Oladele DA, Awoderu OB, Afocha EE, Lawal RG, Abdus-Salam I, Ogunsola FT, Audu RA. 2020. A national survey of public awareness of antimicrobial resistance in Nigeria. Antimicrob Resist Infect Control 9:72.

44. San Millan A. 2018. Evolution of Plasmid-Mediated Antibiotic Resistance in the Clinical Context. Trends Microbiol 26:978–985.

45. Baty D, Knibiehler M, Verheij H, Pattus F, Shire D, Bernadac A, Lazdunski C. 1987. Site-directed mutagenesis of the COOH-terminal region of colicin A: effect on secretion and voltage-dependent channel activity. Proc Natl Acad Sci U S A 84:1152–1156.

46. Marković KG, Grujović MŽ, Koraćević MG, Nikodijević DD, Milutinović MG, Semedo-Lemsaddek T, Djilas MD. 2022. Colicins and Microcins Produced by Enterobacteriaceae: Characterization, Mode of Action, and Putative Applications. Int J Environ Res Public Health 19.

47. Kucerova E, Clifton SW, Xia X-Q, Long F, Porwollik S, Fulton L, Fronick C, Minx P, Kyung K, Warren W, Fulton R, Feng D, Wollam A, Shah N, Bhonagiri V, Nash WE, Hallsworth-Pepin K, Wilson RK, McClelland M, Forsythe SJ. 2010. Genome sequence of Cronobacter sakazakii BAA-894 and comparative genomic hybridization analysis with other Cronobacter species. PLoS One 5:e9556.

48. Franke S, Grass G, Rensing C, Nies DH. 2003. Molecular analysis of the copper-transporting efflux system CusCFBA of Escherichia coli. J Bacteriol 185:3804–3812.

49. Llor C, Bjerrum L. 2014. Antimicrobial resistance: risk associated with antibiotic overuse and initiatives to reduce the problem. Ther Adv Drug Saf 5:229–241.

50. Ecker L, Olarte L, Vilchez G, Ochoa TJ, Amemiya I, Gil AI, Lanata CF. 2011. Physicians’ responsibility for antibiotic use in infants from periurban Lima, Peru. Rev Panam Salud Publica 30:574–579.

51. Adisa R, Orherhe OM, Fakeye TO. 2018. Evaluation of antibiotic prescriptions and use in under-five children in Ibadan, SouthWestern Nigeria. Afr Health Sci 18:1189–1201.

52. Duong QA, Pittet LF, Curtis N, Zimmermann P. 2022. Antibiotic exposure and adverse long-term health outcomes in children: A systematic review and meta-analysis. J Infect 85:213–300.

